# High-throughput multi-camera array microscope platform for automated 3D behavioral analysis of swimming zebrafish larvae

**DOI:** 10.1101/2025.07.07.661868

**Authors:** Haitao Chen, Kevin Li, Lucas Kreiss, Paul Reamey, Lain X. Pierce, Ralph Zhang, Ricardo Da Luz, Amey Chaware, Kanghyun Kim, Clare B. Cook, Xi Yang, Joshua F. Lerner, Jed Doman, Aurélien Bègue, John Efromson, Mark Harfouche, Gregor Horstmeyer, Matthew N. McCarroll, Roarke Horstmeyer

## Abstract

Understanding the behavioral and morphological dynamics of moving model organisms like the zebrafish larvae requires accurate, high-throughput 3D analysis. However, traditional single-view 2D video tracking fails to capture the full scope of natural 3D movements and postural dynamics. Here, we present a novel high-throughput 24-camera array microscope with a co-designed “mirrored well plate” that allows for snapshot imaging of up to 48 wells over a 118 mm ***×*** 82 mm field of view from two orthogonal directions (i.e., a top-view and side-view). Accurate 3D position estimation and tracking is achieved with an efficient machine learning algorithm that scales well to high-throughput measurements. The proposed approach automates parallelized 3D model organism behavioral analysis, providing 3D skeletal tracking, swim bladder morphological dynamics, and kinematics of up to 48 swimming zebrafish larvae at up to several hundred frames per second. The result is an efficient and scalable solution for high-throughput 3D behavioral studies with broad compatibility with standard workflows across laboratories and procedures working with pharmacology, toxicology, and neuroscience.

## 1 Introduction

Behavioral and morphological analysis of zebrafish provides crucial insights into neurological [1], pharmacological [2], and toxicological processes [3], bridging the gap between molecular mechanisms and observable phenotypes [4]. In early studies, morphological and behavioral analyses of zebrafish were usually performed using video tracking from two-dimensional (2D) single-view imaging [5, 6]. However, this setup inevitably restricted their movement and reduced their three-dimensional (3D) trajectories and morphology to a 2D projection [7, 8], preventing the full representation of their natural habits. Despite efforts to estimate the 3D tilt angle of zebrafish based on their head orientation from 2D video [9–11], the resolution of overlapping trajectories of multiple zebrafish [7, 8, 12] and the quantification of behavioral parameters such as speed, acceleration, and total travel distance, among other metrics, remain challenging [12, 13].

By incorporating a second image viewpoint, either through multi-camera stereo vision techniques [7, 8, 13] or monocular stereo using mirrors [12, 14, 15], researchers can resolve these ambiguities and gain access to more accurate kinematic and morphological measurements. This additional view reveals critical 3D postural dynamics, axial behaviors, and physiological responses that often indicate neural or muscular function [16, 17], but are obscured in single-view setups [14, 18]. However, precise orthogonal alignment of multiple cameras is challenging [14]. Moreover, both multi-camera and monocular stereo systems that incorporate a second side view are mainly used to track adult zebrafish in groups at macroscopic scales, where multiple fish share the same tank [8]. These designs offer limited resolution and thus have limited applicability to the study of smaller model organisms such as the zebrafish larvae and Drosophila, especially for analysis of finer details, such as 3D pose estimation or organ segmentation. Other setups can image individual fish at higher resolution and from orthogonal views, but severely limit the free-motion space of the specimen and/or the experimental throughput [19]. All of the above setups are not directly compatible with high-throughput studies, which typically study dozens to hundreds of individual organisms in parallel within multi-well plates.

Current approaches to multi-view batch imaging of zebrafish larvae use microfluidics [20], stepper motors [21], or engineered well plates (funnel or bump) [22]. Although effective, these techniques rely on serial imaging, where each larva is rotated and imaged individually from different angles [23]. They are also time-consuming (up to a few minutes per well plate) and not suitable for high-throughput applications [23]. For example, in the context of drug screening, where acute pharmacological effects on behavior or physiology are critical, the inability to image all animals simultaneously may introduce temporal confounds [24, 25]. This limitation is further compounded when analyzing stimulus-evoked behavioral responses, where precise timing and synchronization across individuals are essential for accurate interpretation. Outside of this field, recent advances in high-throughput multi-view imaging have incorporated the use of mirrors within well plates, which has been used in fast light-sheet 3D imaging of organoid cultures with a single objective [26]. Alternatively, multi-camera array microscopes (MCAMs) [27–31], which use densely packed arrays of micro-cameras to image in parallel and then estimate depth through stereo imaging with overlapping views [32], have been developed for 3D reconstruction of free-moving organisms such as harvester ants and fruit flies [9, 33, 34]. Despite these advances, challenges remain in visualizing 3D information in real time, as it is often derived from computational reconstructions performed offline.

In this paper, we present a novel, custom-designed mirrored multi-well plate in combination with a high-throughput MCAM system for parallel 3D behavioral analysis of swimming zebrafish larvae over an 82 mm *×* 118 mm field of view. Our systems high resolution over this large field of view enables the finer details of zebrafish larvae to be resolved, with the larvae itself free from anesthesia and mounting and only constrained by the physical well walls. The well plate is designed such that half of the wells of a standard 96 well plate are replaced with 45° mirrors. Coupled with an appropriately calibrated MCAM system containing 24 individual high-resolution cameras, we synchronously capture top and side views across all wells (Fig. 1a,b; Video 1). Combined with algorithms for multi-view video tracking and analysis, our platform supports parallelized automated 3D skeletal tracking, 3D morphology measurements (e.g., of the swim bladder) and kinematic quantification of 48 specimens at once (Fig. 1c,d; Video 1). These capabilities make our proposed system highly suitable for high-throughput experiments, which are important in drug discovery, toxicology, and developmental biology, for example. We applied this system to perform 3D analysis of general behavior and response to ethanol exposure in zebrafish, revealing loss of key kinematic details in conventional single view 2D analysis. We have also demonstrated its application in psychoactive drug screening, dissecting drug impact on behavioral dynamics across swim depth. Our system provides a high-throughput, user-friendly approach to studying 3D zebrafish larvae behavior in parallel, and ensuring accessibility and adaptability to standard laboratory workflows and a variety of experimental setups.

**Fig. 1.**
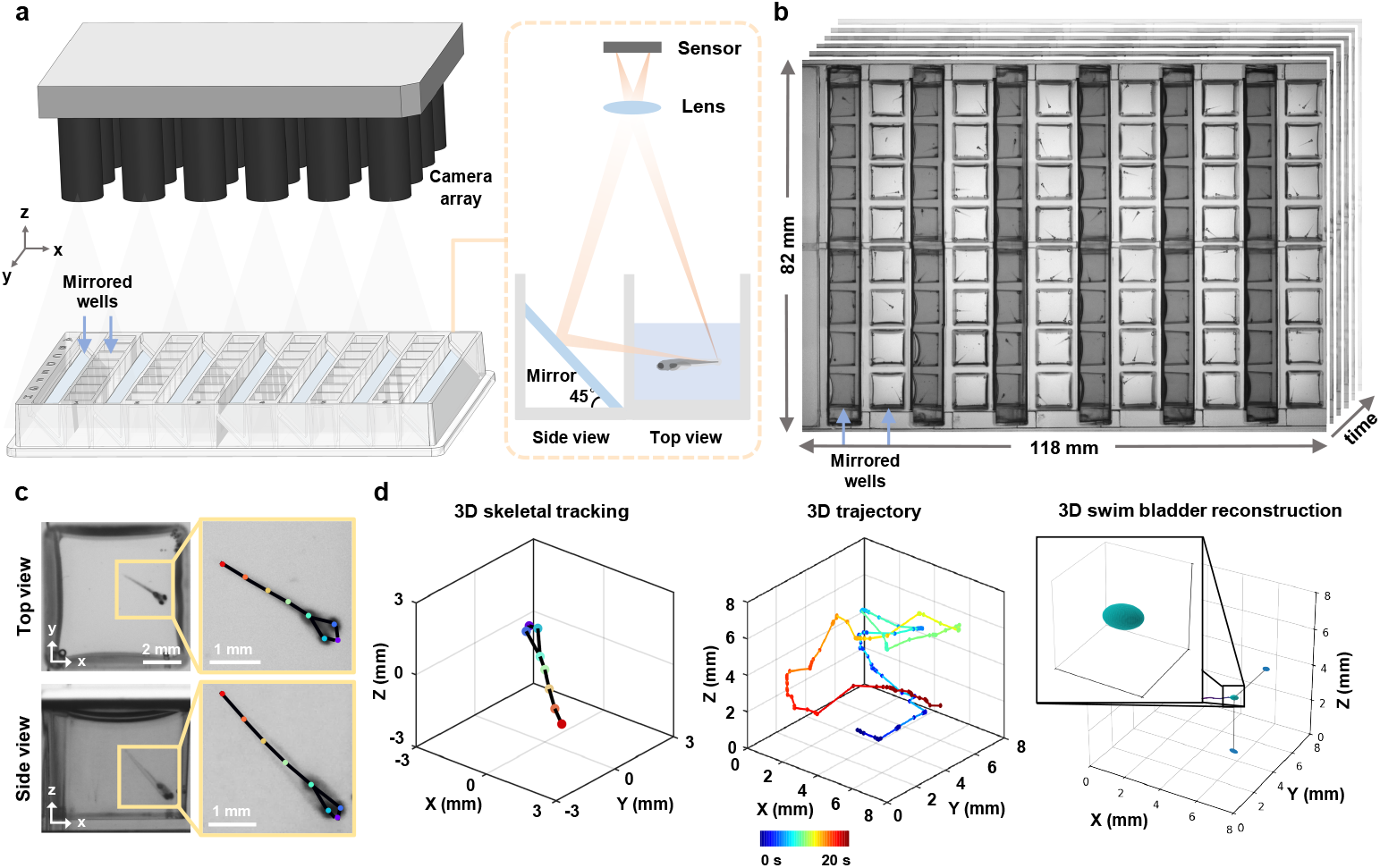
High-throughput multi-camera array microscope platform for parallel 3D behavioral analysis. **a**, 3D rendering of the multi-camera array microscope platform and plug-and-play mirrored well plate with integrated 45° mirrors. **b**, Snapshot dual-view imaging of 48 wells over a 118 mm *×* 82 mm field of view. **c**, Zoom-in views of a mirror well with skeletal tracking. **d**, Automated 3D skeletal tracking (left), 3D trajectory (middle), and 3D swim bladder morphological dynamics (right). Color bar: time.

## 2. Results

### 2.1 High-throughput 3D microscope platform

To enable large-scale 3D behavioral monitoring, we developed a high-throughput microscope platform integrating a customized mirrored well plate (Ramona Optics Inc.) and a MCAM system (earlier prototype of a now commercially available Kestrel system, Ramona Optics Inc.) with 24 micro-cameras arranged in a 6 *×* 4 grid (Fig. 1; Video 1). The custom mirrored well plate conforms to the ANSI/SLAS standard 96 well plate format but replaces half of the wells with mirrors placed at 45° angles, enabling simultaneous acquisition of two orthogonal views (Fig. 1a). Each well has a footprint of 8 *×* 8 mm^2^ and a depth of 12 mm, with a center-to-center spacing of 9 mm between adjacent wells. This standardized design gives it a plug-and-play nature and ensures broad compatibility with standard well plate workflows across laboratories and procedures. The mirrored well plate can be easily integrated with other single-camera imaging systems for ANSI-standard plates, as well as with microplate-based video tracking systems, such as DanioVision. However, additional software development would be required. Depending on whether the MCAM is upright or inverted, our platform allows simultaneous top and side (or bottom and side) imaging of up to 48 wells over an 82 mm *×* 118 mm field of view in a single shot (Fig. 1b).

Each micro-camera features a customized lens (*f* = 25.05 mm, Edmund Optics) and captures up to 3,120 *×* 4,208 pixels with a Complementary Metal Oxide Semi-conductor sensor (CMOS; AR1335, ONSemi) [27], and is arranged on a single Printed Circuit Board (PCB) with a pitch of 18 mm. This configuration provides a 28.5 mm *×* 28.5 mm field of view (FOV) at the 240 mm object distance for each camera, resulting in around 40% overlap in the sample plane FOV between adjacent cameras. A total of ~315 megapixels per snapshot are transferred to computer memory at ~5GB/s via Peripheral Component Interconnect express (PCIe). The LED array panel provides uniform infrared illumination at 850 nm and deep tissue penetration, while ensuring imaging in visually dark conditions (i.e. invisible to the fish eye). This setup allows continuous monitoring of fast, 3D animal motion with minimal behavioral disturbance.

### 2.2 System characterization

The mirror introduces an additional optical path length, which results in slight differences in focal positions of top and side view. To balance the image quality between these two views, we positioned the focal plane in an intermediate position during the imaging (Fig. 2a), leading to a varying magnification between 0.1129 and 0.1205 across the array according to the object distances, which was carefully taken into account when calculating the image pixel size. Given the 9 mm well spacing and 18 mm camera pitch, each camera captures four full wells in the center of its FOV, with partial wells at the edges covered by the central regions of adjacent cameras to enable accurate stitching of composite images (Fig. 1b).

**Fig. 2.**
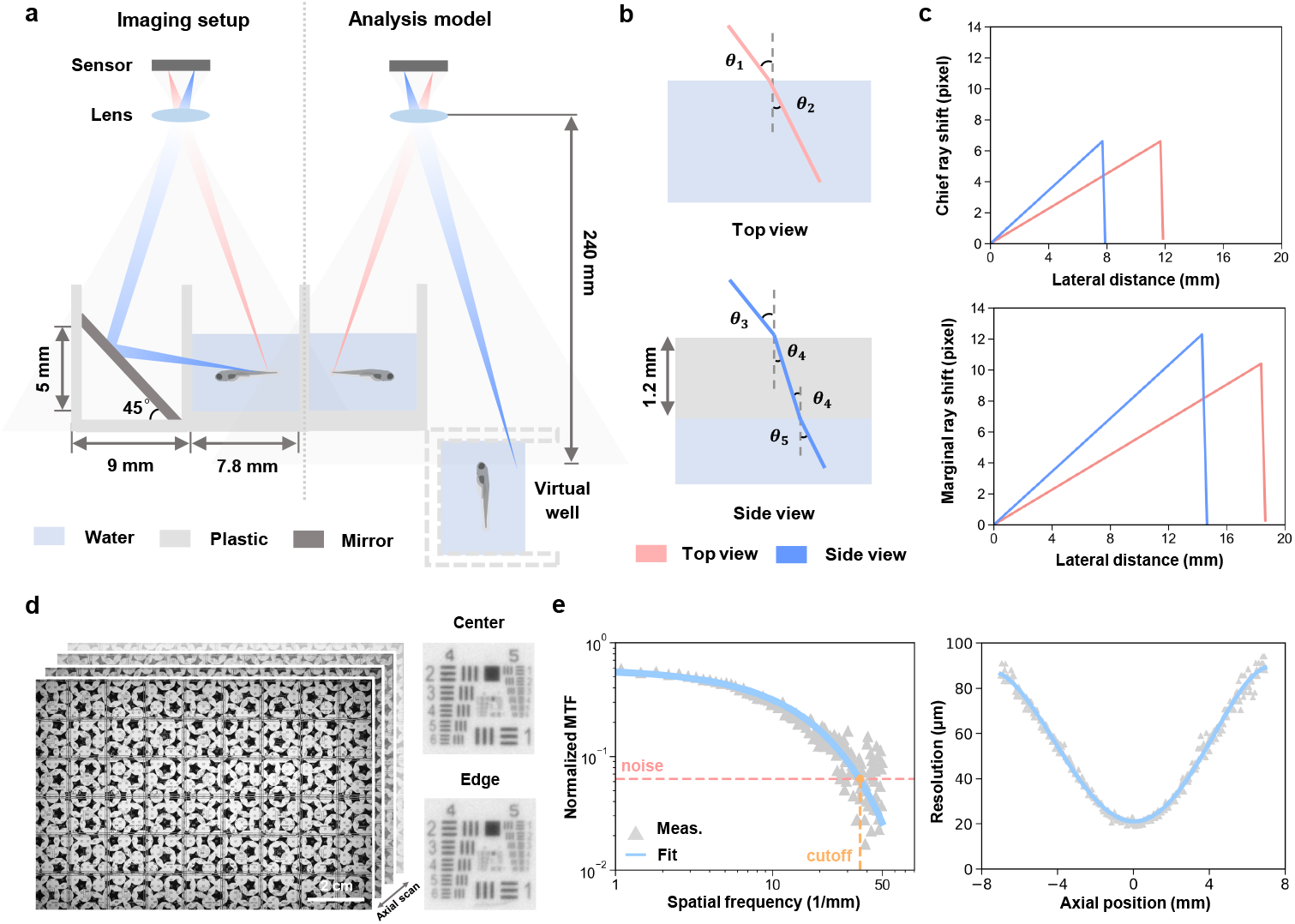
Optical setup and characterization of MCAM. **a**, Schematic of the imaging principle of the multi-camera array microscope platform (left) and its analysis model for a real well and its virtual counterpart by mirror reflection (right). Grey shading represents the field of view of a single camera. **b**, Optical path schematic for top and side view imaging, showing refracted rays passing through air, plastic, and water, with angles of refraction (*θ*_1_–*θ*_5_). **c**, Chief ray (top) and marginal ray (side) distortion corresponding to lateral distance in top and side views. **d**, Axial stack of a resolution target, scanned at 0.1 mm intervals from −7 mm to 7 mm relative to the focal plane. **e**, Modulation transfer function derived from vertical edges in the axial stack (left). The cut-off spatial frequency, determined by the intersection of the fitted exponential decay with the noise floor, was used to calculate the lateral resolution as a function of axial position (right).

Therefore, the analysis of refractive distortions caused by water and the plastic wall were quantified using an equivalent model consisting of only four wells — two real and two virtual ones (Fig. 2a). To assess optical performance, we constructed an equivalent ray-tracing model using Snell’s law, incorporating air-water (*n* = 1.33) and air-plastic (*n* = 1.5) transitions (Fig. 2b, Methods Sec. 4.1). Taking advantage of our relatively long working distance and the small aperture of each camera lens, pixel shifts caused by refraction were small: *<*8 pixels at chief ray’s incident angle (*<* 3° top, *<* 2° side), and *<* 13 pixels in worst-case marginal rays (*<* 5°), leading to an overall distortion *<* 1% (Fig. 2c). Additionally, the short focal length of the lens leads to a magnification of about 0.1, further minimizing any pixel shift caused by refractive distortion.

Lateral resolution variation was assessed using an axial stack of a custom-designed resolution target scanned from −7 mm to +7 mm (Fig. 2d). The small aperture of each camera lens effectively minimizes the variation in resolution across different regions within the FOV of a single camera, consistent with our previous results using the same resolution target [27]. Modulation transfer function (MTF) curves were derived from the vertical edges, fitted to an exponential decay and used to extract the lateral resolution at each axial position (Fig. 2e, Methods Sec. 4.1). Despite defocus at depth, resolution remained sufficient for behavior analysis presented in this work, degrading gradually from 20 µm to 90 µm (Fig. 2e).

### 2.3 Automated 3D skeletal tracking

By integrating depth information from side view imaging, we can closely monitor all degrees of freedom in 3D motion. Building on this, we implemented a deep-learning-based pipeline to track 3D skeletal movements of zebrafish larvae from dual-view videos, providing precise details of fine and fast movements. The overall procedure consists of two primary steps: image pre-processing and skeletal tracking (Fig. 3a, Methods Sec. 4.2). We first enhance and normalize the raw images, ensuring that the data quality is suitable for subsequent analysis. We then perform skeletal tracking to identify and track eight key points of interest (nostrils, eyes, swim bladder and four tail points) and fuse the dual-view data to generate 3D skeletons and perform detailed motion analysis.

**Fig. 3.**
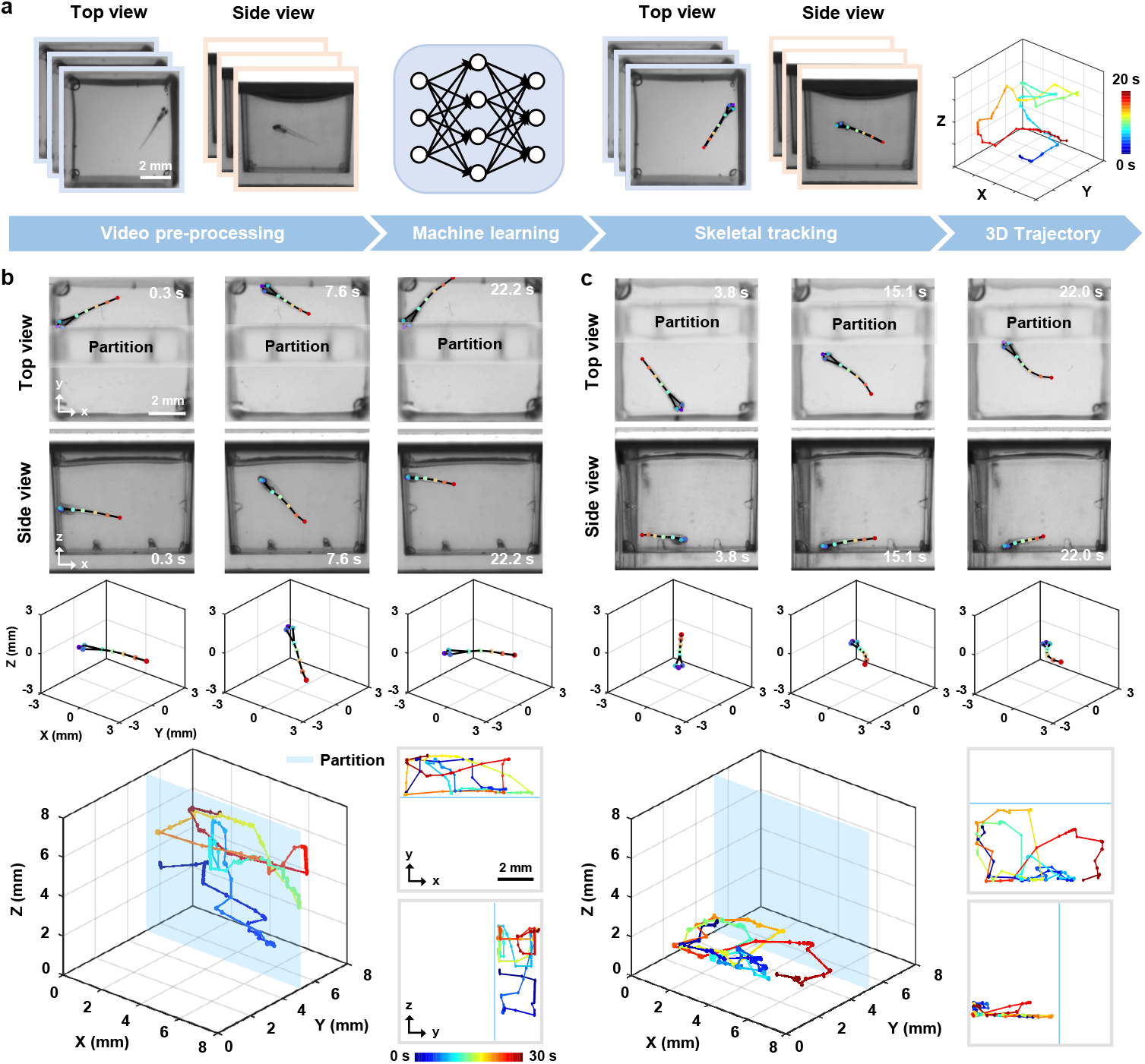
Automated 3D skeletal tracking for motion trajectory analysis. **a**, Workflow of the 3D skeletal tracking pipeline, including image pre-processing and key point extraction of zebrafish larvae using machine learning algorithms. The extracted key points form the skeleton in top and side views and are then combined to reconstruct 3D trajectories. Color bar: time. **b-c**, Skeletal tracking from top and side views at several time points, fused 3D skeletons, and 3D motion trajectories of confined zebrafish larvae (bottom left), and 2D projections along the *x* –*y* and *y* –*z* planes (bottom right).

**Fig. 4.**
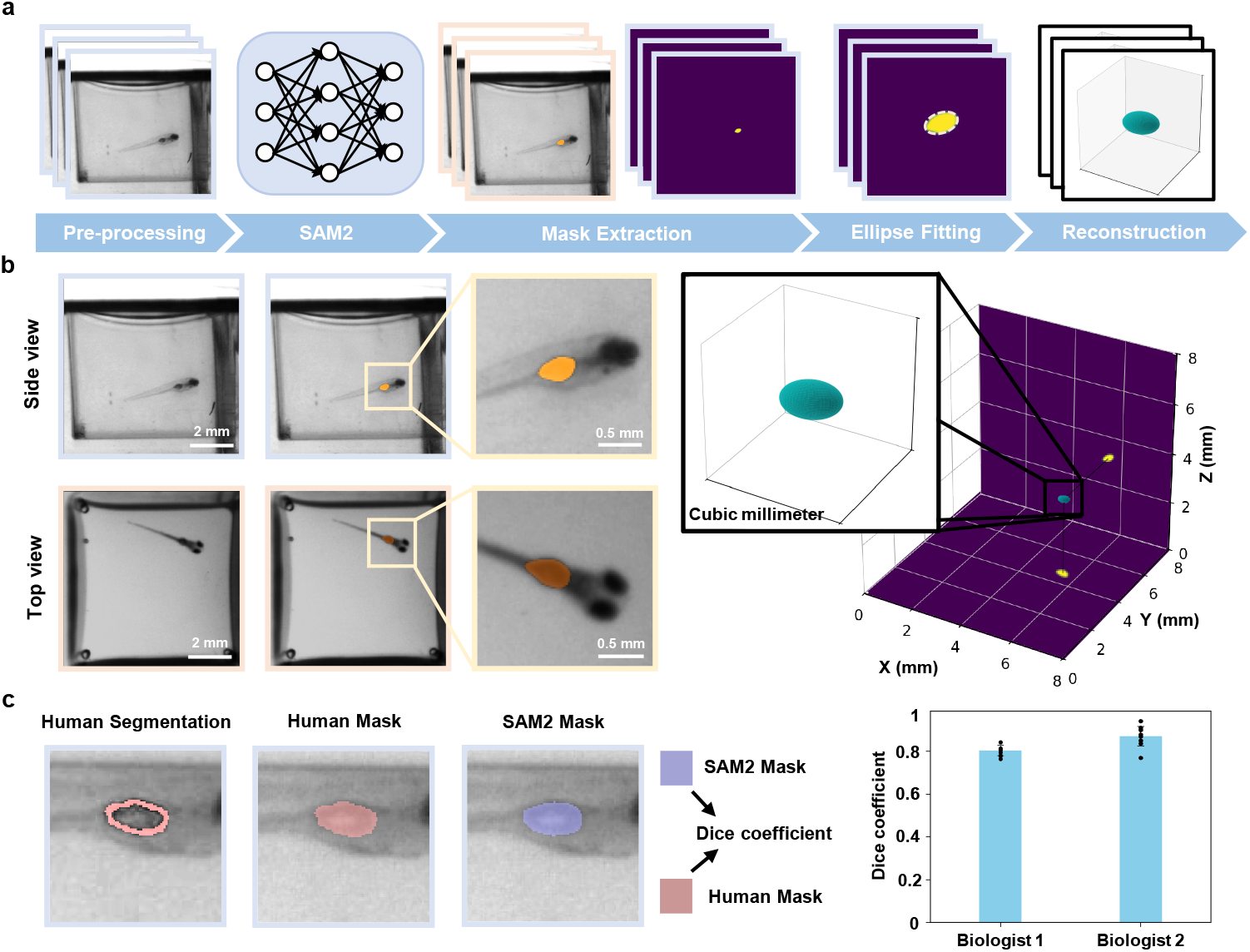
3D swim bladder reconstruction for morphological dynamics analysis. **a**, Workflow of the 3D swim bladder reconstruction. Segmentation is performed using SAM2 on both top and side views. Ellipses are fitted to the segmented contours in each view, which are then used to reconstruct the 3D geometry of the swim bladder. **b**, Visualizations of the swim bladder segmentation and the resulting 3D reconstruction (right). **c**, Representative comparison of segmentation masks generated by SAM2 and independent expert biologists for *n* = 30 zebrafish. Dice coefficients: Biologist 1: mean = 0.80, SD = 0.02; Biologist 2: mean = 0.87, SD = 0.05.

To evaluate the 3D tracking accuracy of our pipeline, root mean square error (RMSE), mean average precision (mAP), and mean average recall (mAR) were calculated on the test set (Methods Sec. 4.2). For the top view, the model achieved an RMSE of 1.94 pixels, mAP of 98.33%, and mAR of 99.49%. For the side view, the RMSE was 2.68 pixels, with mAP and mAR of 96.38% and 96.15%, respectively. These results confirm high prediction accuracy in both views, enabling reliable 3D fusion. Moreover, to further test the robustness and qualitatively validate the plausibility of our pipeline, we restricted the swimming space of the zebrafish larvae by inserting a transparent plastic partition. The partition created a known spatial constraint, allowing us to visually confirm that the reconstructed 3D trajectories remained within the expected boundaries. We then used the MCAM to capture simultaneous top and side views of larvae in 48 wells at 10 frames per second (fps) and performed skeletal tracking using the same model.

Representative tracking results and fused 3D skeletons for two zebrafish larvae are summarized in Fig. 3b,c. The 3D trajectories, along with the *x* –*y* and *y* –*z* projections, confirm that the larvae’s movements were confined entirely to the designated side of the partition. More importantly, our tracking method demonstrated strong robustness across diverse wells and movement types. While gross tail flicks were reliably captured (Fig. 3c), the rapid motor events, such as C-start responses, which occur within ~30 ms [35, 36], cannot be fully resolved at the system’s full resolution. Nonetheless, our approach remains well-suited for capturing the overall morphology and directionality of such behaviors. All imaging was performed with an exposure time of 2 ms, which, while not increasing temporal resolution, ensures that individual frames are free from motion blur, preserving the instantaneous morphology of fast movements. Higher frame rates (up to 160 Hz) are achievable via pixel binning, offering a potential tradeoff between temporal resolution and spatial detail. Furthermore, the consistency of the results tracked across multiple wells demonstrates its reproducibility, with successful 3D skeleton reconstructions achieved in 45 out of 48 wells (93.8%), and a per-frame keypoint tracking success rate of 94.1%*±*2.3% (likelihood *>* 0.7 across all 8 keypoints).

### 2.4 Automated 3D swim bladder morphological dynamics

To showcase the system’s morphological tracking capabilities, we performed 3D swim bladder reconstruction by combining top and side view segmentations of the projected swim bladder (Methods Sec. 4.3). This reconstruction then allows for an accurate estimation of the swim bladder volume for various treatments and conditions. Limitations arise with these initial segmentations when the fish is out of focus, or when orientation is closely facing either of the two views, as the swimbladder cannot be accurately identified behind the eyes of the fish (Fig. S1). We address the latter through a visibility criteria, in which we obtain the true length of the fish via the aforementioned skeletal tracking. We then compare this length to the 2D projected lengths of the fish measured from top and side views, and exclude any frames in which the 2D length of the fish is insignificant relative to the true 3D length, as this suggests that the fish is facing a camera; in our case, we excluded frames where the ratio fell below 0.5.

To validate our 2D mask segmentation of the swim bladder, we compared masks segmented by SAM2 and those by humans. Two biologists were presented with random images of the swim bladder that had passed the visibility criteria, and the areas they segmented were quantified against the masks segmented by SAM2. Using the Dice coefficient [37, 38], we calculate scores of 0.80 and 0.87 with respective variances of 0.02 and 0.05. The low variances suggest that our method is consistent throughout the zebrafish videos.

### 2.5 2D versus 3D kinematic analysis under ethanol exposure

High-throughput, 3D kinematic analysis is a key advantage of our system. To reveal potential information loss inherent in single view 2D kinematic analysis, we selected 16 zebrafish larvae and placed them individually in the mirrored well plate. Of these larvae, eight were treated with 0.5% ethanol (EtOH) to induce acute ethanol responses and stimulate activity, while the remaining eight served as control group. Prior to MCAM acquisition, the larvae were allowed a 5 minute adaptation period in the mirrored well plate. A 30 second video was then recorded at 20 fps to capture the 3D swimming behavior of the larvae.

Following video acquisition, we applied a custom 3D skeletal tracking algorithm to identify and track key feature points of each zebrafish larva. We then quantified both 2D (using top view only) and 3D (combining the top and side views) kinematic parameters, including swimming speed, acceleration, and tail angle. The actograms illustrate the kinematic differences between 2D and 3D tracking for one representative larva from the control group (0.0% EtOH) and one from the group exposed to 0.5% EtOH (Fig. 5a,b). In these comparisons, the 3D actograms revealed consistently greater magnitudes of speed, acceleration, and tail angle than their 2D projections, demonstrating that conventional single-view tracking systematically underestimates core kinematic parameters. More importantly, 3D tracking uncovered behavioral features that were either muted or entirely obscured in 2D analyses, such as nuanced variations in tail curvature and changes in swimming trajectory depth. These finer-grained measures provide access to a richer, more physiologically relevant characterization of locomotor output. For example, larvae exposed to 0.5% EtOH exhibited significantly elevated average speed, increased movement frequency, and exaggerated tail angles during bouts of swimming—behaviors that are likely indicative of acute ethanol-induced perturbations of central nervous system function, consistent with previous reports [39, 40]. These findings highlight the limitations of top-down 2D imaging alone and underscore the value of 3D behavioral phenotyping in revealing subtle drug-induced locomotor changes that may otherwise go undetected.

**Fig. 5.**
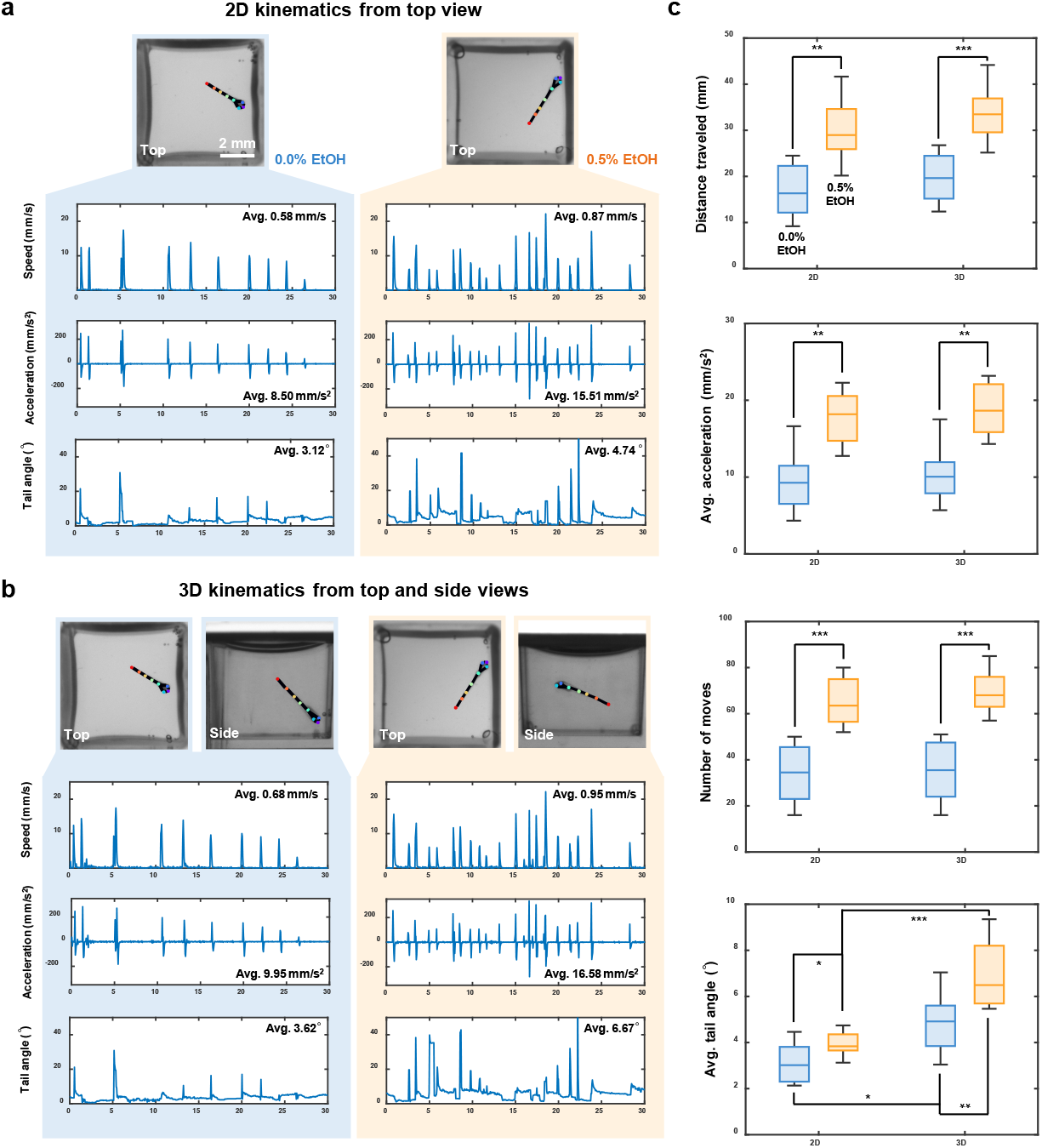
2D and 3D kinematics of zebrafish larvae under ethanol exposure. **a**, Representative 2D tracking actograms for control (left) and ethanol-exposed (right) zebrafish larvae. **b**, Representative 3D tracking actograms for control (left) and ethanol-exposed (right) zebrafish larvae. **c**, Quantitative statistical analysis of kinematic parameters, including distance traveled, average acceleration, number of moves, and average tail angle, between 2D and 3D tracking methods (*n* = 8 per group). p-values were computed using a Mann-Whitney U test. ^*^ *p <* 0.05, ^**^ *p <* 0.01, ^***^ *p <* 0.001. Comparisons without statistical annotations are not significant.

Quantitative statistical analyses of the kinematic parameters for all larvae are shown in Fig. 5c and Fig. S2a, in the form of a box plot and a Nightingale plot [41], respectively. Across all metrics, 2D tracking consistently underestimated movement magnitude compared to the 3D analysis. This systematic underestimate is particularly problematic for long-term or high-throughput behavioral studies, where small errors can compound and lead to misclassification or loss of meaningful biological signal. Among all metrics, tail angle showed the greatest relative benefit from the inclusion of side view data, due to its sensitivity to both lateral and vertical tail bending. This parameter is critical for identifying stereotyped swimming behaviors such as fast-starts or escape responses. As shown in Fig. S2b,c, the larvae exhibit distinctly different bending behaviors when viewed solely from above or from the side. This complex out-of-plane bending can lead to misinterpretations of their swimming behavior. Because tail angle influences hydrodynamic force generation and 3D torque, underestimating this parameter can obscure key biomechanical features of motor output [42, 43]. Importantly, by incorporating a side view into our mirror well plate system, we gain literal access to a previously unmeasured dimension of behavior. This added resolution not only improves the accuracy of existing kinematic measurements, but also opens the door to more precise behavioral classification and the detection of subtle phenotypes that would be missed using top-view-only approaches. In the context of drug screening or genetic perturbation studies, where nuanced shifts in movement patterns can be biologically meaningful, the ability to fully reconstruct 3D behavior greatly enhances the sensitivity and interpretability of phenotypic profiling.

### 2.6 Axial behavior and swim bladder dynamics under neuroactive compounds

The development and screening of drugs for neurological and psychiatric disorders (such as depression, neurodegeneration, and drug addiction) is one of the greatest challenges in modern medicine. In this context, behavioral analysis of zebrafish larvae has emerged as a valuable, non-invasive method to identify new neuroactive substances [24, 25, 44]. In some contexts, their response to neuroactive compounds is closely linked to swim bladder homeostasis and swimming depth regulation [45, 46]. However, to thoroughly dissect the intrinsic neural mechanisms underlying buoyancy control, batches of data derived from numerous individual larvae are required. High-throughput, automated axial behavior analysis and morphological dynamics are other distinct advantages of our system. Leveraging these capabilities, we performed behavioral imaging and 3D analysis of zebrafish larvae exposed to two neuroactive compounds and vehicle: 25 µM 3-CPMT, 12.5 µM Nicergoline, and Dimethyl sulfoxide (DMSO). These compounds act on distinct neuropharmacological targets—3-CPMT as a dopamine uptake inhibitor and Nicergoline as an *α*-adrenergic and serotonergic antagonist with pro-cognitive effects—both of which are known to influence motor control, arousal, and sensorimotor integration. Following a 15-minute exposure, 30-second dual-view recordings were acquired at 20 fps and analyzed using our 3D skeletal tracking and organ segmentation pipeline. Capturing axial depth and swim bladder morphology in swimming larvae is particularly innovative, as these parameters have traditionally been inaccessible in high-throughput paradigms. Axial positioning (i.e., vertical displacement) is closely tied to vestibular, buoyancy, and sensorimotor function, while swim bladder inflation is a sensitive physiological readout of neural control and internal pressure regulation. By enabling automated, high-resolution assessment of these previously underutilized dimensions, our platform increases the sensitivity of behavioral profiling and opens the door to detecting subtle, multidimensional phenotypes that may be missed with conventional 2D analysis.

Top view behavior appeared similar across groups; however, side views revealed distinct axial motion profiles (Fig. 6a). The difference is further illustrated through 3D trajectories and heatmaps of a representative larva from each treatment group compared to the control group in Fig. 6b. Control larvae displayed more distributed axial activity, and their swim bladder volume lays evenly distributed over the height axis (Fig. 6e). In contrast, larvae exposed to Nicergoline reduced the larvae’s range of motion and restricted their mobility near to the water surface (Fig. 6b,c), where they exhibited larger swim bladders. This reduced mobility is probably due to vasodilation, which limits the oxygen supply to the swim bladder, resulting in marked bladder inflation, and high buoyancy [47]. Larvae treated with 3-CPMT exhibit the opposite, remaining in contact with the bottom of the well for almost the entire time, with occasional upward swims followed by rapid sinking (Fig. 6b,c). This behavioral pattern is in agreement with previous reports [48], indicating the sensitivity of buoyancy regulation to neuroactive compounds and demonstrating that exposure to these drugs disrupts swim bladder homeostasis. The cluster of low volume deflated swim bladders are reflected in the volume measurements. These results are consistent with the findings of Venuto et al. [19] who recently demonstrated the correlated relationship between buoyancy and the volume of zebrafish swim bladders. However, while Venuto et al. used a conventional stereo microscope for imaging and manual bladder segmentation in Adobe Photoshop, our approach massively increases experimental throughput by exploiting parallelized imaging with the MCAM and AI-supported bladder segmentation.

**Fig. 6.**
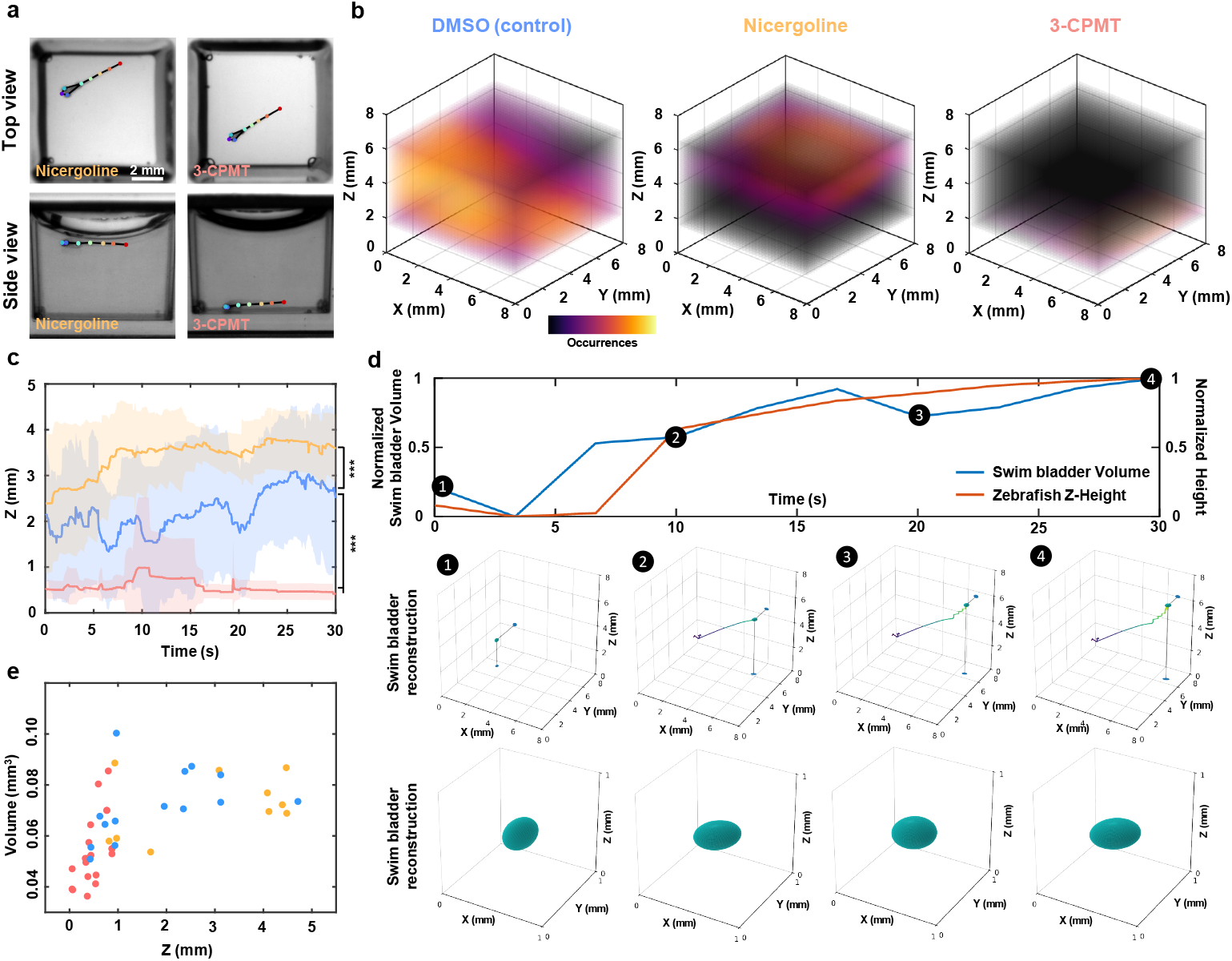
Axial behavior and swim bladder dynamics under neuroactive compounds. **a**, Representative top view (upper row) and side view (bottom row) images of zebrafish larvae treated with Nicergoline (left column) and 3-CPMT (right column). Colored dots indicate tracked anatomical landmarks. **b**, 3D occupancy heatmaps showing typical swimming behaviors for larvae treated with DMSO (control, left), Nicergoline (center), and 3-CPMT (right). **c**, Quantitative analysis of swimming depth over time for larvae exposed to DMSO (blue), Nicergoline (orange), and 3-CPMT (red). Solid lines denote mean z position, and shaded areas indicate standard deviation (*n* = 8 per group). p-values were computed using a Mann-Whitney U test. ^***^ *p <* 0.001. **d**, Representative positive correlation between bladder volume and zebrafish height within the well. 4 time points are extracted, each displaying the swim bladder location and 3D reconstruction. **e**, Swim bladder volume height correlation graph for the three experimental groups.

By capturing both locomotor dynamics and internal morphological features—such as swim bladder volume and axial posture—our platform provides a more holistic and mechanistically informative readout of compound effects. This added dimensionality improves behavioral resolution, enhances the detection of subtle drug-induced phenotypes, and facilitates classification of neuroactive compounds based on complex movement and physiological signatures. Together, these advances position our system as a next-generation tool for probing neurobehavioral function with both depth and precision. Therefore, our system holds promise as a powerful high-throughput 3D behavioral screening tool that can uncover phenotypes missed by conventional 2D assays and enable large-scale 3D behavioral studies to accelerate psychoactive drug discovery.

## 3 Discussion

In this work, we presented a novel high-throughput multi-camera array microscope platform, incorporating a plug-and-play mirrored well plate that enables snapshot top and side imaging of swimming zebrafish larvae across 48 wells over an 82 mm *×* 118 mm field of view. The combination of multi-view imaging and machine learning algorithms automates 3D skeletal tracking, kinematics and swim bladder morphological dynamics, revealing key kinematic and morphological details (such as tilt angles and axial movement) that are lost in top view setups.

Our approach has several practical advantages over conventional methods. First, the mirrored well plate retains the standard 96-well plate format, facilitating straightforward integration into existing laboratory workflows. Second, simultaneous multi-view imaging eliminates the need for time-consuming sample rotation or serial image acquisition. Third, parallel data acquisition and automated skeletal tracking speed up the collection and analysis of large datasets of individual larvae, improving reproducibility and scalability. With these combined benefits, our plug-and-play, high-throughput, multi-camera array microscope platform allows comprehensive 3D behavioral studies to be performed on a large scale, providing an invaluable tool for a wide range of disciplines, from pharmacological testing and toxicological screening to neuroscience and behavioral ecology.

Our analysis of 3-CPMT—a known dopamine transporter (DAT) inhibitor—revealed distinctive 3D behavioral alterations, including persistent bottom-dwelling, that closely resemble phenotypes reported in adult zebrafish with loss-of-function mutations in the DAT gene (slc6a3) [49]. Previous work has shown that DAT-deficient zebrafish exhibit hypolocomotion and altered vertical positioning, often swimming near the bottom of the tank, suggestive of impaired dopaminergic regulation of motor control and arousal states. The similarity between this pharmacologically induced phenotype and its genetic counterpart highlights the ability of our 3D platform to capture behaviorally meaningful patterns that reflect underlying neurochemical mechanisms. Importantly, such depth-resolved behaviors are rarely detectable using traditional 2D imaging approaches. By accessing this additional spatial dimension and combining it with detailed skeletal tracking, our system expands the repertoire of accessible behavioral phenotypes, enabling the dissection of pharmacological effects, genetic perturbations, and their interactions with neural circuit function. This positions the platform not only as a tool for psychoactive compound screening, but also as a versatile framework for studying the behavioral architecture of neurological function and disease.

Looking beyond zebrafish, the flexible design of our platform opens up a wide range of possibilities. From aquatic animals to mollusks and even invertebrates such as tardigrades [50], our platform enables 3D locomotor and morphological analyses that can be adapted to the specific needs of different model organisms and experimental paradigms. Similarly, organoids [51] or other 3D cell cultures grown in specialized multi-well formats can benefit from simultaneous multi-angle imaging to track growth, morphological changes and functional markers over time.

A limitation of the system was its low depth of field, which prevented successful segmentation of the fish swim bladder at certain positions of the well, even to human observers. In the future, this could be mitigated by the integration of different lenses, or by reducing the size of the wells. Additional hardware and computational upgrades that could further extend the functionality of the platform could include the integration of multiple spectral channels to use fluorescent markers or genetically encoded indicators for insights into neural activity, muscle contraction, or cell signaling events. In addition, the use of higher magnification autofocus lenses, where each lens captures only one view of the well, could further refine the balance between optimizing resolution for both the top and side views. Furthermore, employing more sophisticated machine learning techniques may facilitate deeper insights into group dynamics and complex behaviors across a wider range of species.

To promote adoption and stimulate continued development, we present this work as an open invitation to the research community. We encourage others to implement, adapt, and build upon the platform described here. The system is composed of three modular and accessible components: the imaging hardware, the mirrored well plate, and the image analysis software. Although the multi-camera array was developed in collaboration with Ramona Optics, comparable configurations can be assembled using commercially available individual cameras. Detailed specifications for the mirrored plate are provided to facilitate replication, and the full image analysis pipeline is openly available via GitHub. This accessibility enables researchers across disciplines to apply, customize, and extend the platform for their specific needs. By emphasizing modularity and openness, we aim to support broad applicability and foster community-driven innovation in imaging-based biological research.

## 4 Methods

### 4.1 Resolution and distortion analysis

To assess lateral resolution, we used a translation stage to obtain an axial stack of a custom-designed resolution target to evaluate the lateral resolution. Images were captured at 0.1 mm intervals across a range of −7 mm to 7 mm. A clear vertical edge was manually selected from each axial stack. We applied a spatial frequency response algorithm to derive the modulation transfer function (MTF), as described earlier [31], plotting the MTF across spatial frequencies from 0 to 50 cycles per millimeter. The MTF is divided into two regions: an exponential decay region and a noise cut-off region at higher spatial frequencies. The exponential decay region was fitted with an exponential function and the cut-off spatial frequency was defined as the intersection of this fitted curve with the mean noise floor. The reciprocal of this cut-off frequency was then used as a measure of the lateral resolution for each axial position.

The analysis of refractive distortions introduced by water and the plastic wall was based on only four wells — two real and two virtual ones. Starting at the optical axis of the camera, we virtually scanned potential chief rays in increments of 0.1 mm until reaching the farthest point from the optical axis in either the real or virtual well. This process produced a series of air-water incident angles (*θ*_1_) or air-plastic incident angles (*θ*_4_) (Fig. 2b). Snell’s law was applied to account for refractive index mismatches, using 1.33 for water and 1.5 for plastic. Rays traveling through water were tracked until they reached a boundary, at which point they were terminated. Shifts relative to air imaging were calculated, and the worst-case scenario was considered, where a marginal ray produced the largest incident angle in air (Fig. 2b).

### 4.2 3D skeletal tracking and kinematic quantification

We developed a Python-based dual-view 3D skeletal tracking pipeline consisting of image pre-processing and pose estimation.

In the first step, dual-view images captured by the MCAM images were cropped to extract views of 48 wells, downsampled to 512 *×* 512 pixels, rescaled to brightness values between 0 and 255, and saved as individual MP4 videos. In the second step, three random frames per view were selected from each well and clustered using k-means to reduce similarity. These frames were then manually annotated with eight key points: the nostrils, both eyes, the swim bladder, and four points along the tail. We reserved 15% of the annotated data as a held-out test set and evaluated model performance using standard metrics: root mean square error (RMSE), mean average precision (mAP), and mean average recall (mAR). We trained separate HRNet-w32 networks via DeepLabCut [52] for top and side views (300 epochs) on an NVIDIA RTX 2080 Ti GPU. Finally, only frames with network prediction likelihood *>*0.7 were retained, and entire video was excluded if over one-third of its frames were below this threshold. Valid data yielded precise kinematic quantification including 3D displacement, velocity, acceleration, and tail angles based on dynamic fused 3D skeletal reconstruction from top and side views.

Midpoint of the line connecting the two eyes of the zebrafish larvae was chosen as the 3D reference point, as the head is the only rigid part of the fish’s body. The Euclidean distance between this reference point in two consecutive frames is calculated to measure the displacement between frames. Based on this displacement and the imaging frame rate, we derive the speed and acceleration. Finally, by summing the inter-frame displacements, the total travel distance is obtained. In addition, the line connecting the nostril to the reference point is considered the forward vector of the zebrafish larvae, and the angle between this vector and the line connecting the nostril to the tail is defined as the tail angle.

### 4.3 Swim bladder segmentation

Our application involves segmenting out the swim bladder of the fish in both top and side views, before modeling a 3D volume reconstruction.

For that purpose, we use Segment Anything Model 2 (SAM2) to initially segment the swim bladder in both top and side view videos [53]. We can operate under the reasonable assumption that the swim bladder follows an ellipsoidal shape, as previously done in similar approaches [19]. We then fit both segmented swim bladders to a 2D elliptical fit. These 2D ellipses are then projected into a 3D ellipsoid under a least-square optimization. Upon converging to the closest ellipsoid solution, a magnification factor based on the height of the fish within the well is applied to obtain the final swim bladder volume.

The matrix form of the ellipse is represented as *u*^*T*^ *Q*^*T*^ *AQu* = 1, where *u* is the position vector, *A* is the matrix containing the major and minor axes, and *Q* is the matrix describing the rotation:

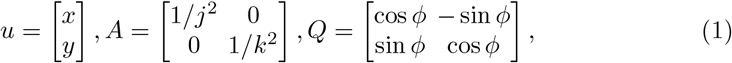

where *j, k*, and *ϕ* are all determined by the fitting function fitEllipse() from OpenCV.

The matrix form of the ellipsoid similarly can written as : *v*^*T*^ *R*^*T*^ *BRv* = 1 where *v* is the position vector, *B* is the matrix containing the major and minor axes, and *R* is the matrix describing the rotation:

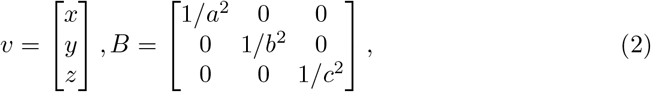

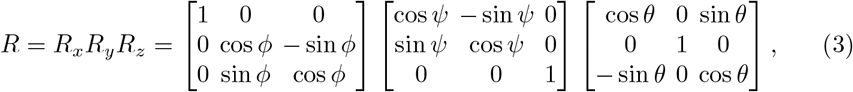

where *a, b, c* are the major and minor axes of the ellipsoid. *ϕ, ψ*, and *θ* are the rotations about the axes *R*_*x*_, *R*_*y*_, *R*_*z*_

The variables *a, b, c, ϕ, ψ, θ* are then least-squares optimized to minimize the error between the projections of the 3D ellipsoid and the previously fitted 2D ellipsoid:

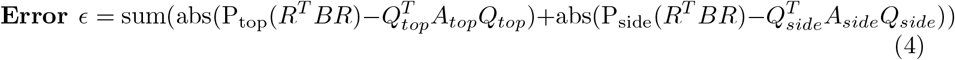

### 4.4 Fish maintenance and breeding

Fertilized zebrafish embryos were obtained from group matings of wild-type zebrafish (Singapore strain). Larvae were maintained on a 14:10 hour light/dark cycle at 28°C and raised until 7 days post-fertilization (dpf). Prior to behavioral testing, larvae were anesthetized in chilled egg water and transferred into custom-built mirrored well plates, with 1 larvae per well. Plates were incubated at room temperature for 30 minutes to allow recovery from cold exposure and resumption of spontaneous activity.

### 4.5 Compounds and treatments

All compounds were prepared as 30 mM stock solutions in Dimethyl sulfoxide (DMSO) and further diluted in egg water to achieve the desired final treatment concentration. Ethanol (EtOH) was diluted directly in egg water. DMSO-treated controls received equivalent vehicle volumes. For compound retesting, treatments were replicated across 8 wells. Drug solutions were added directly to the egg water, and larvae were incubated for durations ranging from 5 to 15 minutes prior to behavioral analysis, depending on the time course being studied.

All zebrafish procedures were performed in accordance with protocols approved by the University of California, San Francisco (UCSF) Institutional Animal Care and Use Committee (IACUC), and in compliance with the Guide for the Care and Use of Laboratory Animals (National Institutes of Health, 1996).

## Data availability

The data supporting the results of this study are available in the paper and its supplementary information. The raw data are too large to be publicly shared, but they are available for research purposes from the corresponding authors upon reasonable request.

## Code availability

The code used for skeletal tracking was based on DeepLabCut (https://github.com/DeepLabCut/DeepLabCut), an open source toolbox for markerless pose estimation. For swim bladder segmentation, we used the publicly available SAM2 (https://github.com/facebookresearch/sam2) repository. All custom code used in this study will be made available on GitHub upon publication.

## Acknowledgements

Research reported in this publication was supported by the Office of Research Infrastructure Programs (ORIP), Office of the Director, National Institutes Of Health of the National Institutes Of Health and the National Institute Of Environmental Health Sciences (NIEHS) of the National Institutes of Health under Award Numbers R44OD024879 and R44OD036187, the National Cancer Institute (NCI) of the National Institutes of Health under Award Numbers R44CA285197 and R44CA250877, the National Institute of Mental Health (NIMH) of the National Institutes of Health under Award Number R43MH133521, the National Institute of Biomedical Imaging and Bioengineering (NIBIB) of the National Institutes of Health under Award Number R43EB030979, the National Science Foundation under Award Number 2036439, the Duke-Coulter Translational Partnership Grant, NIDA DP1DA058350, P0569108 NIH (Ramona SBIR), and Sandler Program for Breakthrough Biomedical Research (PBBR) project 7031733.

## Contributions

H.C., K.L., G.H., and P.R. conceived the idea. H.C. performed optical performance measurements and 3D skeletal tracking. K.L. conducted swim bladder segmentation and 3D reconstruction. H.C., K.L., L.K., A.C., K.K., C.B.C., X.Y., and J.L. processed visualizations. M.H., G.H., P.R., J.D., A.B., J.E., and R.H. developed the mirrored well plate, MCAM hardware, and acquisition software. L.X.P., R.Z., R.D.L., and M.N.M. acquired and analyzed the biological data. M.N.M. provided input to and supervision of the biological experiments. H.C. and K.L. wrote the manuscript with input from all authors. H.C., K.L., L.K., L.X.P., R.D.L., M.N.M., and R.H. revised the manuscript. L.K., M.N.M., and R.H. supervised the research.

## Supplementary information

### S1 Supplementary video

1. **System principles, 3D skeletal tracking, and swim bladder morphodynamics**.

### S2 Additional figures

**Fig. S1.**
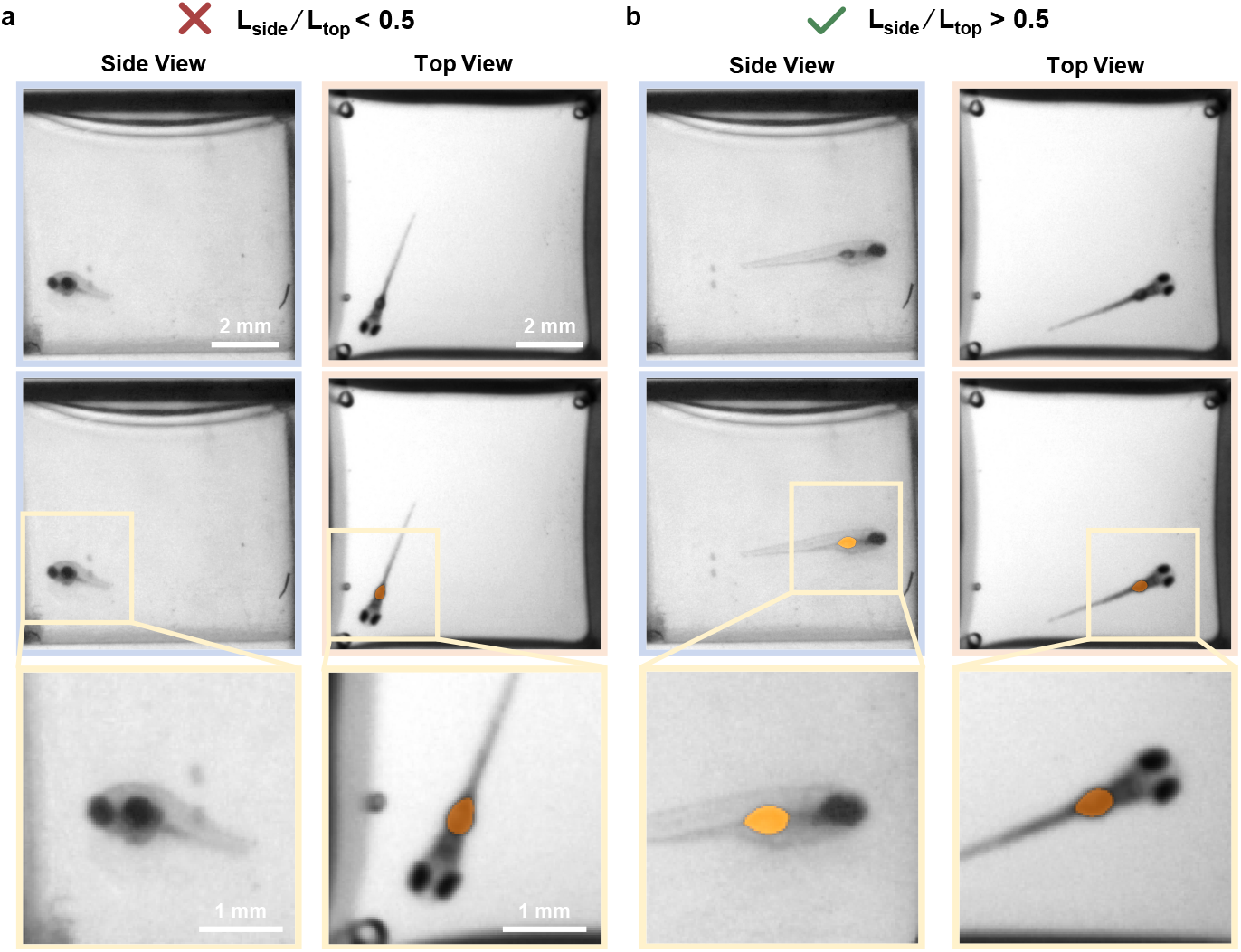
Representative examples of our visibility criteria for segmenting the swimbladder. **a**, Visibility criteria is not met, the fishes projected length on the sideview is short relative to the true length, resulting in the swimbladder being behind the zebrafish’s eyes. **b**, Visibility criteria is met, the long projected lengths in both views ensure that the swim bladder is clearly visible and segmentable.

**Fig. S2.**
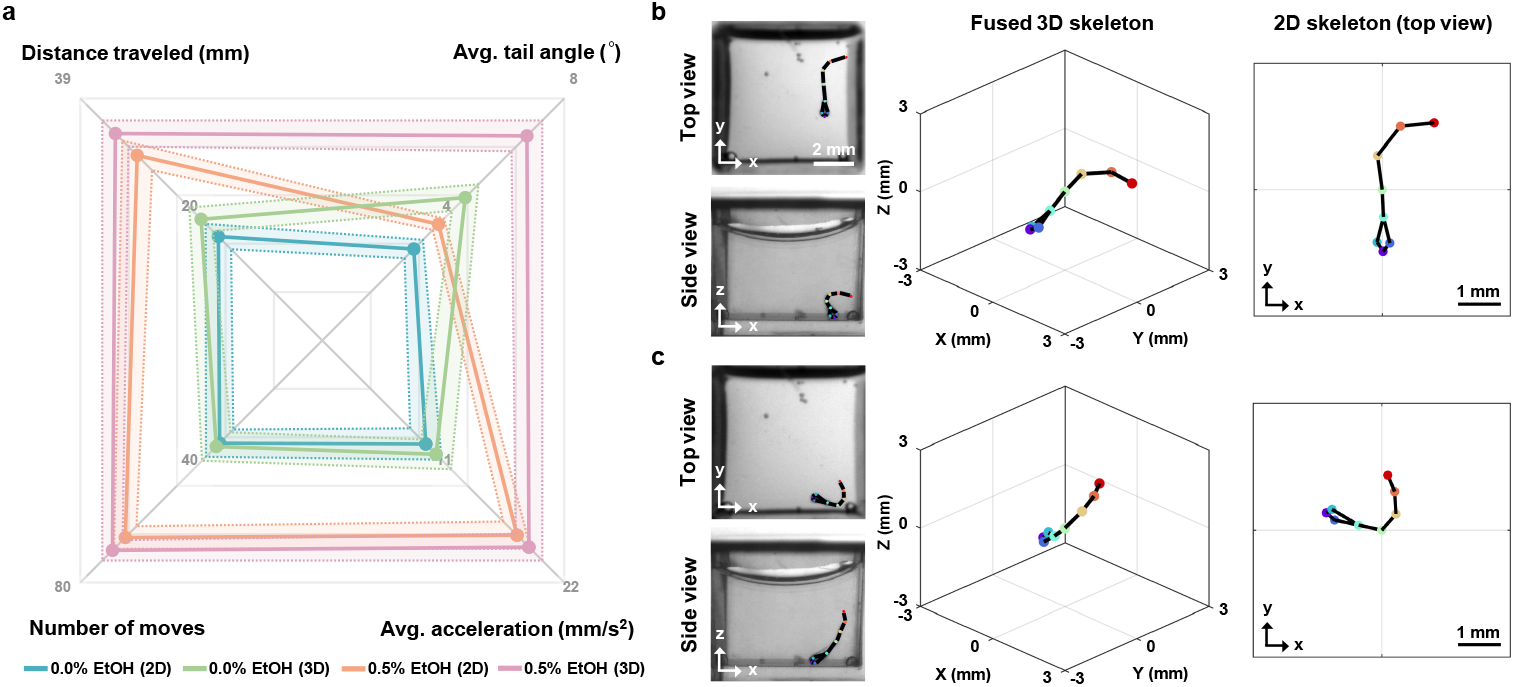
Comparison of 2D and 3D kinematic analysis under ethanol exposure using multivariate visualization and skeletal reconstruction. **a**, Nightingale plot illustrating the average values and standard errors (SEM, shaded regions) of four key kinematic parameters for zebrafish larvae exposed to 0.0% or 0.5% ethanol (EtOH), measured using 2D and 3D tracking (*n* = 8 per group). **b-c**, Two examples of larvae exposed to 0.5% EtOH. In **b**, the top view suggests a classic J-turn, while the side view reveals that the movement is actually a C-turn with strong vertical curvature. In **c**, the side view reveals an upward arched tail bend not captured from above.

